# Chewing behavior and bolus particle size of rice shape gut microbiota functionality and microbial metabolite signatures

**DOI:** 10.1101/2025.03.07.642000

**Authors:** Zhen Liu, Ciarán G. Forde, Markus Stieger, Josep Rubert

## Abstract

The size of food particles and their surface area affect how accessible the substrates are to the gut microbiota. This study aimed to examine how the oral breakdown of rice influences the gut microbiota functionality, considering the oral phase, digestion, and colonic fermentation. Increasing the number of chews per bite produced more and smaller particles and increased total bolus surface area, influencing microbial activity and microbial metabolite production. Higher bile salt hydrolase activity and total short-chain fatty acid (SCFA) levels were found in large particles. Focusing on SCFAs, acetic acid was correlated with larger particle sizes. Propionic acid production was associated with a larger surface area, while butyric acid production was highly correlated to the fiber content. Untargeted lipidomics highlighted that particle size influenced numerous metabolites beyond SCFA. This study showed that food particle size could be an innovative strategy to modulate gut microbiota functionality and microbial metabolite production.

## 1. Introduction

The texture of the food and the characteristics of an individual’s preferred chewing behavior determine the properties of the bolus at swallow (Bornhorst & Singh, 2012). Oral processing behavior varies considerably across people (Chen et al., 2021; Tang et al., 2025). During oral processing, food matrices are minced into smaller particles and lubricated with saliva to form a bolus that is subsequently swallowed. More chews per bite increase the number and decrease the size of bolus particles, resulting in an increased total surface area of the bolus. The bolus particle size after chewing can affect gastric digestion and gastric emptying (Nadia et al., 2021a). Bolus particles with smaller sizes have larger surface areas and are more easily broken down by digestive enzymes, leading to increased nutrient absorption (Teo et al., 2022), glucose release, and insulin production (Goh, Choy, et al., 2021). However, whether these differences in bolus particle size affect digestion in the upper and lower intestines and the functionality of microbial communities remain unknown.

In the upper and lower intestines, bolus particles’ size and surface area may be crucial in regulating the gut microbiome and its metabolites. Indeed, the metabolic activity of the gut microbiome in the colon is considered an essential part of food digestion (Letourneau et al., 2024), since the gut microbiome utilizes nutrients and influences intestinal homeostasis (Fernandes et al., 2023; Ravel et al., 2014). For example, before entering the colon, bile salt hydrolase (BSH) catalyzes the deconjugation of bile salts (Zhou & Hylemon, 2014), converting them into free bile acids to be recycled (Wahlström et al., 2016), and indirectly influencing cholesterol metabolism and gut microbiota composition. Emerging evidence indicates that changes in bile acid metabolism may be associated with functional gastrointestinal disorders like irritable bowel syndrome (IBS) (Camilleri, 2015; Vijayvargiya et al., 2018). Another critical group of microbial metabolites is short-chain fatty acids (SCFA), produced by the microbial fermentation of non-digestible carbohydrates in the colon and extensively studied due to their metabolic significance (de Vos et al., 2022; Flint et al., 2012; van Leeuwen et al., 2023). SCFAs are directly involved in histone deacetylase (HDACs) inhibition to regulate gene expression (Koh et al., 2016), protecting the intestinal barrier and controlling inflammation (Deleu et al., 2021). In addition, SCFAs can act as signaling molecules that activate free fatty acid receptors (FFAR), which are G protein-coupled receptors (GPCRs), primarily including FFAR2 and FFAR3 (Kasubuchi et al., 2015; Mohammad, 2015; Rothschild et al., 2018; van der Merwe, 2021).

We hypothesized that bolus particle size and surface area are key abiotic factors impacting microbiome activity and the production of microbial metabolites. Several *in vitro* studies have investigated the impact of particle size on the gut microbiome. Tuncil et al. reported that wheat bran particle size modulated SCFA production during *in vitro* fermentation. While coarse wheat bran produced less acetic and propionic acid, fine wheat bran boosted butyric acid (Tuncil et al., 2018). Other studies have reported the influence of whole oat grain particle size (Connolly et al., 2010) and maize bran (Thakkar et al., 2020) in regulating SCFA production *in vitro*. However, it remains unclear whether bolus particle size influences microbial metabolism and other microbial metabolites beyond SCFAs. Although oral processing behavior plays a crucial role, none of the studies mentioned above have examined the impact of chewing on the microbiome (Koç et al., 2013). Using cows and mice as models, previous studies have shown that a wheat diet’s varying particle size influenced digestibility and gut microbiota composition (Bao et al., 2016; Suriano et al., 2017). More recently, the field has shifted towards considering the entire digestive process, from the mouth to excretion. Kim and co-authors reported that interindividual differences in the chewing behavior of brown rice led to differences in bolus particle size, which influenced colonic fermentation (Kim et al., 2022). In this study, the authors observed that chewing more per bite resulted in smaller bolus particle sizes and increased propionic acid production during colonic fermentation (Kim et al., 2022). In an *in vivo study*, Letourneau and co-authors performed a behavioral intervention with 41 volunteers consuming their habitual diets. They showed that the gut microbiome shaped fecal particle size, affecting metabolite production and transit time (Letourneau et al., 2024). The study reported that mastication did not affect fecal particle size. It is important to note that participants’ chewing behavior and bolus characteristics before and after that intervention were not measured. Nevertheless, this research study emphasized the significance of particle size on the gut microbiome and metabolites.

The current study investigated whether differences in bolus properties induced by differences in human oral processing behavior for rice influence digestion and human microbial metabolism *in vitro*. We hypothesize that (a) increasing the number of chews per bite leads to increased structural breakdown (increased number of smaller bolus particles and increased total bolus surface area) after the oral and gastrointestinal phases and that (b) an increased total surface area increases the activity of microbial enzymes, and therefore, the production of microbial metabolites, such as SCFAs. To test our hypothesis, we use a combination of various *in vivo* and *in vitro* models to minimize the variation from the individual, such as the effects of intestinal length, peristaltic forces, enzymatic concentrations, and transit time, while enabling the tracking of particles from the mouth to the colon. To reduce the impact of dietary heterogeneity, we chose three varieties of rice (white rice, basmati brown rice, and pandan brown rice) differing in fiber content to determine whether the number of chews influences bolus particle size and surface area modulates microbial metabolism. The properties of bolus particles were quantified after a specified number of chews. Subsequently, the oral, gastrointestinal, and colonic phases were correlated with the degree of starch hydrolysis during digestion, SCFA production, BSH, and the pattern of microbial metabolites during colonic fermentation.

## 2. Materials and methods

### 2.1 Materials

Basmati brown rice (B), pandan brown rice (P), and white rice (W) were purchased in a local supermarket (Albert Heijn BV, Wageningen, The Netherlands) and stored at room temperature. The composition of the rice, as shown on the package, is summarized in *Supplementary Table S1*. Following the instructions on the package, the three rice varieties (100 g) were cooked in boiling water (500 mL) for 12-15 min, and the water was then decanted. All rice samples were masticated within 10 min of preparation. Pepsin (P6887, 3200–4500 units/mg protein), pancreatin (P1750, 4 × USP specification), and porcine bile extract (B8631) were purchased from Sigma-Aldrich Ltd. All other chemicals used were analytical grade.

### 2.2 *In vivo* mastication and bolus collection

One participant was instructed for the *in vivo* mastication, which removed inter-individual differences in chewing behavior. The chewing frequency of rice has been reported to range from 1.1 to 1.5 chews/s, so a chewing frequency of 1.25 chews/s was selected for this study (Forde et al., 2013, 2017). A tone was used to prompt a chew by the participant every 0.8 s, corresponding to a chewing frequency of 1.25 chews/s. A fixed bite size of 10 g of rice was chosen based on previous studies (Ekuni et al., 2012; Naumova et al., 2021a). The participant was instructed to chew 10 g of freshly cooked rice 5, 10, 20, and 40 times at a chewing frequency of 1.25 chews/s, corresponding to a chewing duration of 4, 8, 16, and 32 s per bite. Then, the participant expectorated the rice bolus into a petri dish. The participant was provided with water to rinse the mouth between bites. Three boli were collected per rice variety and selected chews. Expectorated boli (10 g) were stored immediately on ice and analyzed within 10 min after expectoration.

Expectorated boli were split for analysis of bolus saliva uptake (3 g), bolus particle properties (1 g), and *in vitro* digestion (5 g). Hereafter, basmati brown rice chewed 5, 10, 20, and 40 times per bite is referred to as B5, B10, B20, and B40, pandan brown rice is referred to as P5, P10, P20, and P40, and white rice is W5, W10, W20, and W40.

### 2.3 Determination of bolus saliva uptake

Saliva uptake of the rice bolus was determined by dry matter content analysis (Devezeaux de Lavergne et al., 2015). Cooked rice bolus samples were collected after different numbers of chews per bite (0, 5, 10, 20, 40 chews) and placed on pre-weighted aluminum dishes and dried overnight in an air oven at 105 °C until constant weight was reached. Three replicates were carried out. The weight of rice bolus and unchewed rice before and after drying was recorded (m_Wet bolus_, m_Dry bolus_, m_Wet rice_, m_Dry ric_e) and saliva uptake was calculated as:

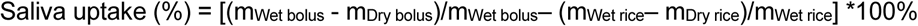

### 2.4 Characterization of bolus particle size, number of particles, and total bolus surface area after mastication, intestinal digestion, and colonic fermentation

Rice bolus properties (particle size, number of particles, total bolus surface area) were determined after the oral (*in vivo* mastication), gastrointestinal (*in vitro* INFOGEST), and colonic fermentation phases using image analysis (Chen et al., 2022). Rice boli (0.5 -1.0 g) were placed in a Petri dish (120×120×17 mm) and weighed. Bolus particles were gently separated using a spatula. After scanning (Canon CanoScan 9000F Mark II) images were analyzed using ImageJ (version 1.52a, National Institute of Health) to obtain particle size (expressed as surface area of each particle), number of particles (expressed as number of particles/g) and total bolus surface area (expressed as total surface area/g). To reduce noise, particles smaller than 0.07 mm2 or with a circularity <0.15 were discarded from data analysis. Particles were categorized into six groups (<0.01, 0.01-0.05, 0.05-0.10, 0.10-0.20, 0.20-0.30, >0.30 cm^2^) to obtain a bolus particle size distribution. For each picture, the number of particles in each size group (-/g), total number of particles (-/g), total bolus surface area (cm^2^/g), and bolus particle mean size (cm^2^) were obtained in triplicate.

### 2.5 *In vitro* gastric and intestinal digestion

Rice bolus (5.0 g) expectorated after different numbers of chews (see section 2.2) were subjected to *in vitro* static digestion immediately after expectoration following the harmonized INFOGEST protocol (Brodkorb et al., 2019). Three replicates were carried out per rice and chewing time. Five g of expectorated rice bolus was mixed with 5 ml Milli-Q water. The mixture was mixed with 6.4 mL prewarmed (37°C) simulated gastric fluid and 5 μL of 0.3 M CaCl2. After changing pH to 3 using 1 M HCl, different volumes of Milli Q water were added to compensate for different volumes of HCl added to the digestion system. Pepsin was added then (final concentration: 2000 U/mL), and gastric digestion at 37°C for 2 h was started. For intestinal digestion, 11 mL of pre-warmed simulated intestinal fluid and 2.5 mL of bile salts were mixed with the gastric digesta. Then, pH was raised to 7 using 1 M NaOH after adding 40 μL of 0.3 M CaCl2 (final concentration:10 mM). Pancreatin was added to reach the final concentration of 200 U/mL for amylase in pancreatin. After 2 h incubation at 37°C, digesta were centrifuged (4500 g, 15 min), and pellets were collected and stored at −20°C after snap freezing in liquid nitrogen. Pellets were further analyzed for bolus properties (section 2.4) and used as a substrate to perform fecal batch fermentation (section 2.7).

### 2.6 Determination of starch hydrolysis

The degree of starch hydrolysis (DH%) was measured before gastric digestion (0 min), after gastric digestion (120 min), after 30 min of intestinal digestion (150 min), and after 120 min of intestinal digestion (240 min) corresponding to the end of intestinal digestion. The degree of starch hydrolysis was analyzed by the quantification of reducing sugar using the 3,5-dinitrosalicylic acid (DNS) assay (Vidal et al., 2009). Briefly, 200 μL of supernatant was taken at each time point during gastric and intestinal digestion. Subsequently, 800 μL of ethanol was mixed thoroughly with the supernatant to stop the reaction. After 30 min at room temperature, the mixture was centrifuged for 10 min at 10,000 rpm and the supernatant was collected. Then, 0.1 mL of ethanolic supernatant was incubated with 0.5 mL amyloglucosidase solution (27 U/mL) in acetate buffer (0.1 M, pH 4.8) for conversion of α-amylase products into glucose at 37°C for 2 h. Samples after incubation were analyzed. DNS solutions were prepared by dissolving 5 g of 3,5- dinitrosalicylic acid and 150 g of sodium potassium tartrate in 500 ml water. Glucose standards with concentrations between 0-1 mg/mL were used to obtain a calibration curve. Samples or standards were mixed with DNS solution in a 1:1 ratio and then incubated at 100°C for 5 min. Reacted samples were cooled to room temperature and diluted using water before measuring absorbance at 540 nm in a microplate spectrophotometer. Glucose content was converted into the corresponding amount of starch by multiplying with a factor of 0.9. The analysis was completed in triplicate for each rice sample bolus across the different numbers of chews.

### 2.7 *In vitro* fecal batch fermentation

*In vitro* fecal batch fermentation was completed following a method described previously with minor modifications (Pérez-Burillo et al., 2021). Fecal samples were provided by four healthy volunteers (20-30 years old). Before donating fecal samples, the volunteers declared no antibiotics or drugs known to influence fecal microbiota consumption for the last 6 months. Healthy volunteers gave written consent for fecal donation, and their anonymity was always maintained. According to the guidelines of the Medical Ethical Advisory Committee of Wageningen University (https://www.radboudumc.nl/en/patient-care), this research (2023-16931) did not require approval by the Medical Ethical Advisory Committee. We performed two *in vitro* fecal batch fermentation studies: (a) the discovery phase consisted of four independent colonic fermentations with intervals of one week using the fecal inoculum of one participant, and (b) the validation phase, in this case, the feces of four donors were included in the experimental design (n=4).

A phosphate buffer consisting of 8.8 g K2HPO4, 6.8 g KH2PO4, and 0.1 g of sodium thioglycolate in 1 L demi water was prepared and sterilized before fermentation. 15 mg sodium dithionite was added to the phosphate buffer before use. Fresh fecal samples within 2 hours of defecation were mixed with the sterilized phosphate buffer in a ratio of 1: 5 (w/v). After mixing using a Stomacher 400 circulator (Seward, UK), the fecal mixture was centrifuged at 500 g for 2 min. The supernatant was collected as fecal inoculum. The undigested pellets collected after *in vitro* gastrointestinal digestion were mixed with fecal inoculum and sterilized basal medium in vessels. Vessels were flushed with N2/CO2 (80/20, v/v) gases to create an anaerobic condition. Samples were taken after fermentation at 0, 4, 8, 12, and 24 hours. Fermentation was stopped by snap-freezing using liquid nitrogen. Samples were stored at −20 °C before analysis.

### 2.8 SCFA analysis during colonic fermentation

Gas chromatography equipped with a flame ionization detector (GC-FID, GC-2014, Shimadzu, Hertogenbosch, Netherlands) was employed to analyze SCFA (acetic acid, propionic acid, butyric acid, isobutyric, isovaleric acid, valeric acid) following the method described previously (Huyan et al., 2022). Nitrogen was used as a carrier gas. The temperature of GC-FID increased from 100 °C to 180 °C at a rate of 10.8 °C min^−1^ maintained at 180 °C for 2 min. Then, the temperature was maintained at 240 °C for 2 min after it increased at 50 °C min^−1^ from 180 to 240 °C. An internal standard of 2-ethylbutyric was added to both fermentation samples and standards. Three replicates were carried out.

### 2.9 Bile salt hydrolase (BSH)

Enzymatic assays were set up to analyze bile salt hydrolase according to a previously described two-step method (Foley et al., 2021). Enzymatic reactions were carried out at 37 °C in 50 μL volumes containing 0.1 M sodium phosphate pH 6.0, 10 mM DTT, 9 mM BA, and 1 to 10 nM BSH. Reactions were quenched using an equal volume of 15% (wt/vol) trichloroacetic acid and were centrifuged at 12,000 × g for 2 min at room temperature to pellet precipitate. To determine the quantity of released amino acid by ninhydrin reaction, 25 μL of the quenched reaction was added to 475 μL of Ninhydrin buffer (0.3 mL glycerol, 0.05 mL of 0.5 M citrate buffer pH 5.5. 0.125 mL of 0.5 M citrate buffer pH 5.5 with 1% [wt/vol] Ninhydrin). The Ninhydrin reaction was developed by boiling for 14 min and cooling on ice for 3 min. A standard curve of glycine or taurine was prepared for each assay. Absorbance at 570 nm was measured in a clear flat-bottom plate in a plate reader.

### 2.10 Untargeted lipidomics

Metabolites were extracted with the method described elsewhere using Ostro 96 well plates (Waters, USA) (Ulaszewska et al., 2019). The untargeted lipidomics determination followed the method described previously using UHPLC (LC-40, Shimadzu, Kyoto, Japan) system equipped with BEH C18 (2.1 × 100 mm, 1.7 µm) analytical column (Huyan et al., 2024). MS-Dial (version 5.52) performed data mining for baseline filter, peak detection, retention alignment, and feature annotation.

### 2.10 Statistical data analysis

Results are shown as mean ± standard deviation (SD). The variance was analyzed using ANOVA followed by Duncan’s post hoc test. Bivariate Spearman correlation (two-tailed) tests were used to examine the relationships between the number of chews, fiber content, rice bolus properties (total surface area, particle size, particle number), degree of starch hydrolysis, and SCFA production. Statistical data analysis, except lipidomics, was performed using IBM SPSS Statistics (version 28.0), and a significance level of *P* < 0.05 was chosen unless stated elsewhere. MetaboAnalyst 6.0 was employed to perform statistical analysis of lipidomics. Before analysis, features with RSD>25% in QC and interquartile range (IQR) < 40% were discarded. Time series + one factor and Multivariate Empirical Bayes Time-series Analysis (MEBA) were used to explore changes in metabolites across different time points of the colonic fermentation.

## 3. Results

### 3.1 Saliva uptake

Saliva uptake of the rice bolus increased significantly with increasing number of chews per bite for the three rice varieties (*Fig. S1*). When the number of chews per bite increased from 5 to 40, saliva uptake of the rice bolus increased 2.2-fold for basmati brown rice, 2.5-fold for pandan brown rice, and 4.9-fold for white rice. Saliva uptake of the rice bolus differed significantly between rice varieties for all numbers of chews per bite, with basmati brown rice showing the highest saliva uptake, followed by pandan brown rice and white rice.

### 3.2 Bolus properties of rice after *in vivo* mastication, *in vitro* digestion, and *in vitro* colonic fermentation

*Fig. 1* and *2* show bolus properties (*Fig. 1*: bolus particle size distribution and the total number of bolus particles; *Fig. 2*: total bolus surface area and average bolus particle size) after *in vivo* mastication, *in vitro* digestion, and *in vitro* colonic fermentation (0, 4, 8, 12, and 24 h). After *in vivo* mastication, the total number of bolus particles increased significantly with increasing chews per bite for all rice varieties. The bolus particles smaller than 0.01 cm^2^ increased by 12.7-fold for basmati brown rice, 8.4-fold for pandan brown rice, and 3.5-fold for white rice when the number of chews per bite increased from 5 to 40. The total bolus surface area increased by 24-47% when the number of chews per bite rose from 5 to 40, and the average bolus particle size decreased by 63-75% for all rice varieties (*Fig. 2*). The reduction of average bolus particles size induced by chewing was more pronounced in basmati and pandan brown rice compared to white rice.

**Fig. 1.**
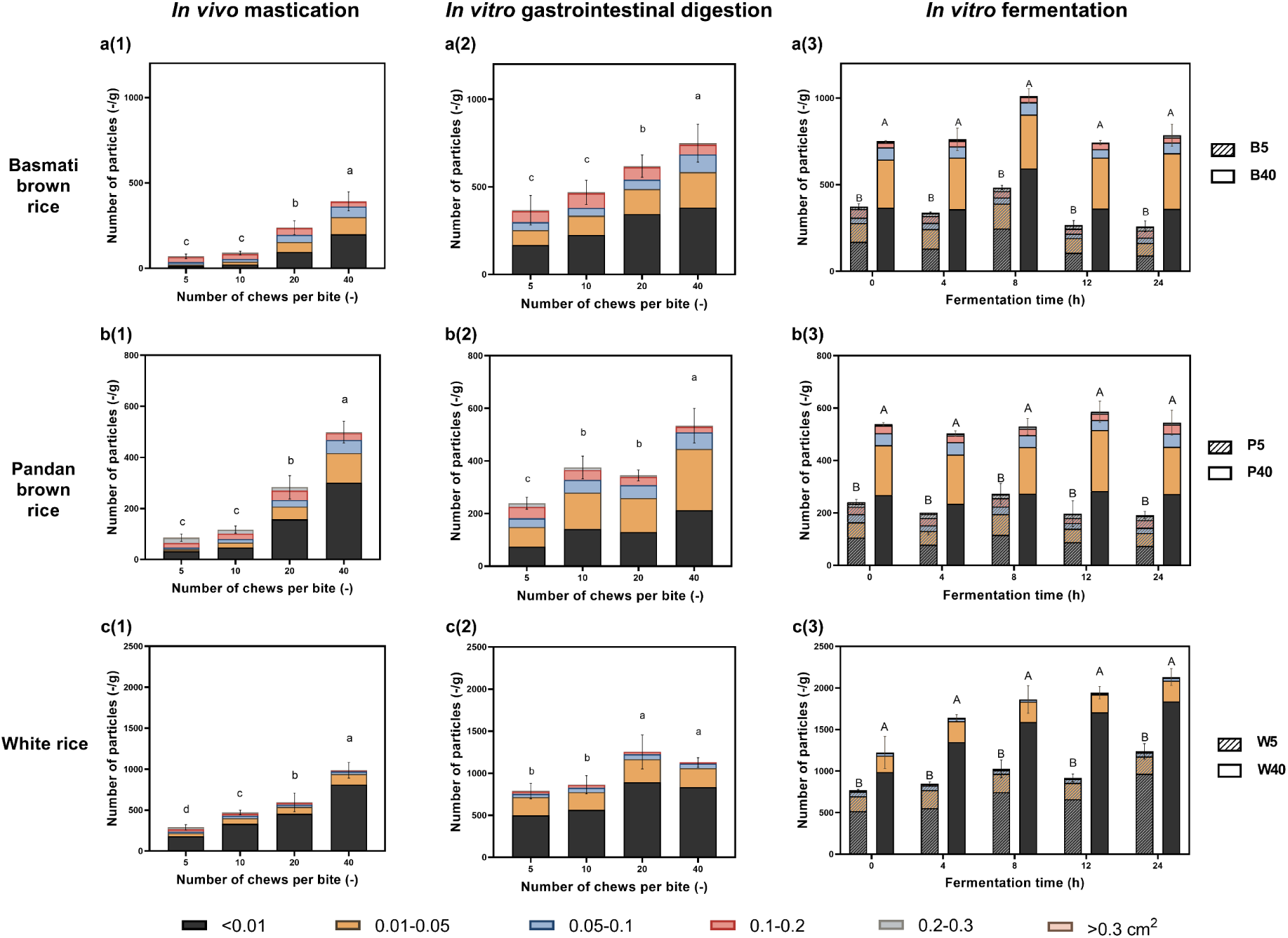
Total number of rice bolus particles (-/g) and number of rice bolus particles (-/g) per size group (<0.01, 0.01-0.05, 0.05-0.10, 0.10-0.20, 0.20-0.30, >0.30 cm^2^) of Basmati brown rice (a), Pandan brown rice (b), and White rice (c), after *in vivo* oral processing (1), after *in vitro* gastrointestinal digestion (2) and after *in vitro* colonic fermentation (3). Stripped and solid bars indicate number of rice bolus particles (-/g) after 5 and 40 chews per bite after *in vitro* colonic fermentation; respectively. Different lowercase letters represent significant differences in total number of rice bolus particles across number of chews per bite after *in vivo* oral processing and *in vitro* gastrointestinal digestion(*P<*0.05). Different uppercase letter denote significant differences in total number of rice bolus particles across number of chews per bite (5 vs. 40) after colonic fermentation for different times.

**Fig. 2.**
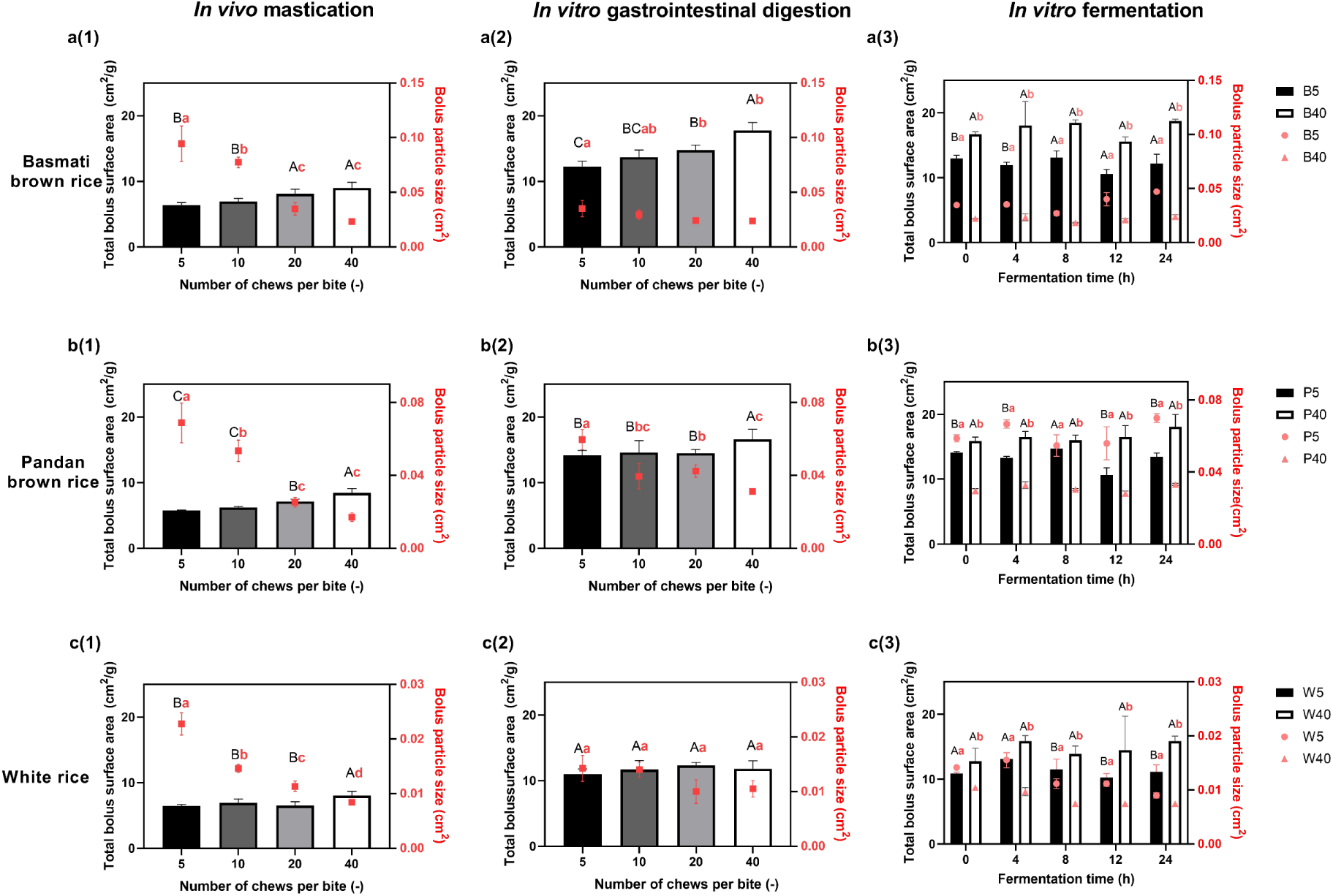
Total bolus surface area (cm^2^/g) and average bolus particle size (cm^2^) of rice after *in vivo* oral processing, *in vitro* gastrointestinal digestion, and *in vitro* colonic fermentation. (a) Basmati brown rice, (b) Pandan brown rice, (c) White rice after *in vivo* oral processing (1), after *in vitro* gastrointestinal digestion (2), and after *in vitro* colonic fermentation (3). The left y-axis (bars) represents total bolus surface area and the right y-axis (red symbol) represents average bolus particle size. Different capital letters represent significant differences in total bolus surface area across number of chews per bite after *in vivo* oral processing and *in vitro* gastrointestinal digestion (*P<*0.05), and different lowercase letters represent significant differences in average bolus particle size across number of chews per bite after *in vivo* oral processing and *in vitro* gastrointestinal digestion (*P<*0.05).

After subjecting the expectorated rice boli to *in vitro* digestion, the effect of oral processing (number of chews per bite) on most bolus properties remained significant. A significantly higher number of bolus particles was observed after *in vitro* digestion for all rice varieties. The average bolus particle size across the rice varieties after the *in vitro* digestion was 26-47% lower after 40 chews per bite compared to 5 chews (*Fig. 1*). Total bolus surface area after *in vitro* digestion rose with increasing number of chews per bite from 10.9-14.2 cm^2^ after 5 chews to 11.8-17.7 cm^2^ after 40 chews across rice varieties (*Fig. 2*). However, the total surface area of white rice after gastrointestinal digestion did not differ significantly (*P*>0.05) across the number of chews per bite. The number of large particles (>0.2 cm^2^) decreased, and the number of small bolus particles (<0.2 cm^2^) increased for all digested rice varieties. The total surface area after *in vitro* digestion significantly increased by 47-147% compared to the total surface area after *in vivo* mastication, suggesting additional structural breakdown of the rice bolus during *in vitro* digestion.

After the *in vitro* gastrointestinal digestion, the undigested rice fractions of 5 chews and 40 chews containing different numbers and sizes of particles were utilized as substrates for the gut microbiome. The effect of oral processing (number of chews per bite) on most simulated fecal properties remained significant during colonic fermentation (*Fig. 1*). A significantly higher total number of particles (1.6-4.2 fold) was found in all rice varieties chewed 40 times compared to 5 times during the colonic fermentation, emphasizing the impact of oral mastication on particle properties during colonic fermentation. The total surface area was significantly higher (8-55%), with a considerably smaller average particle size (17-53%) for all rice varieties after 40 chews compared to 5 chews during the colonic fermentation (*Fig. 2*). Minor differences were observed in the total surface area and the average size of white rice compared to basmati brown rice and pandan rice.

### 3.3 Degree of starch hydrolysis during gastrointestinal digestion

*Fig. 3* shows the degree of starch hydrolysis during the *in vitro* digestion. The degree of starch hydrolysis (DH) significantly increased with digestion time, especially during the intestinal phase of digestion (120-240 min). For the three rice varieties, longer chewing resulted in a significantly higher degree of starch hydrolysis, especially after the oral and gastric phases of digestion (0 and 120 min). Increasing the number of chews per bite from 5 to 40 increased the absolute DH% after the oral phase of digestion (0 min) by 17% (basmati brown rice), 22% (pandan brown rice), and 9% (white rice). After the *in vitro* gastric digestion (120 min), the impact of the number of chews per bite on the degree of starch hydrolysis was more significant than after the oral phase of digestion. Increasing the number of chews per bite from 5 to 40 increased the absolute DH% after the *in vitro* gastric phase of digestion (120 min) by 32% (basmati brown rice), 30% (pandan brown rice), and 26% (white rice). Intense starch hydrolysis occurred during the *in vitro* intestinal phase (150 min). DH% increased by 11-34% (basmati brown rice), 20-44% (pandan brown rice), and 21-40% (white rice) compared to the gastric phase of digestion (120 min). No significant effect of the number of chews per bite (5, 10, and 20 chews) on DH% was observed at the end of the gastrointestinal phase of digestion (240 min) for all three rice varieties. Only the pandan brown rice chewed 40 times showed significantly higher starch hydrolysis than 5, 10, and 20 chews. The white rice chewed 20 times had significantly higher DH% than the other chews at the end of the gastrointestinal phase of digestion (240 min).

**Fig. 3.**
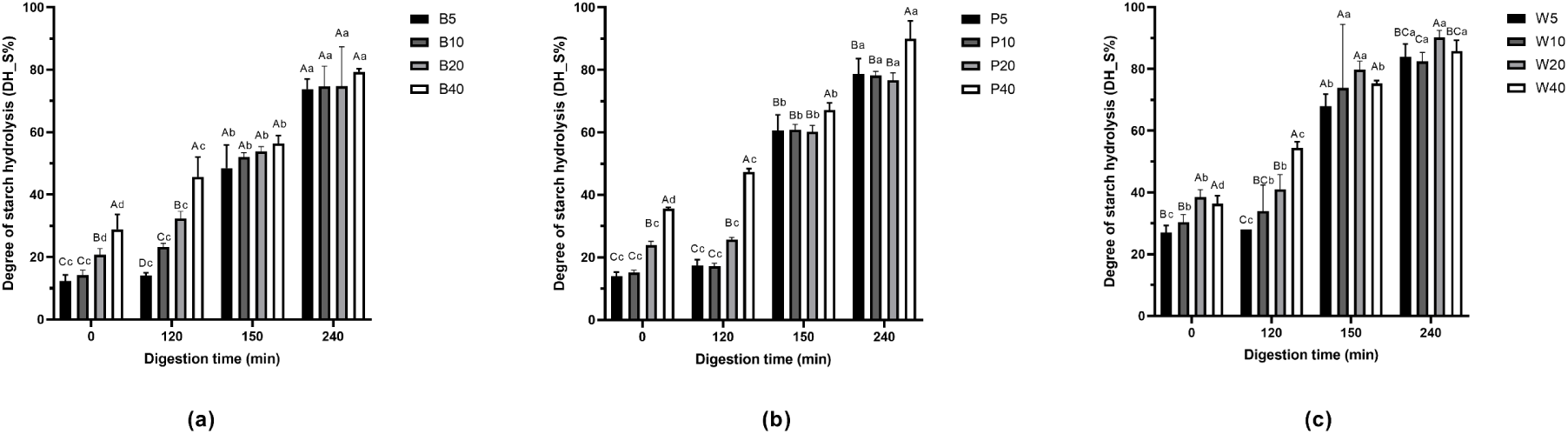
Degree of starch hydrolysis (DH_S%) during gastrointestinal digestion. 0 min: before gastrointestinal digestion (after oral mastication), 120 min: after 120 min *in vitro* gastric digestion, 150 min: after 120 min *in vitro* gastric digestion and 30 min of *in vitro* intestinal digestion, 240 min: after 120 min *in vitro* gastric digestion and 120 min of *in vitro* intestinal digestion. (A) Basmati brown rice, B5, B10, B20, B40 refer to basmati brown rice chewed 5, 10, 20, 40 times per bite, respectively. (B) Pandan brown rice, P5, P10, P20, P40 refers to pandan brown rice chewed 5, 10, 20, and 40 times per bite, respectively (C) White rice, W5, W10, W20, W40 refers to white rice chewed 5, 10, 20 and 40 times per bite, respectively. Different capital letters represent significant differences across numbers of chews per bite at a given digestion time (*P<*0.05), and different lowercase letters represent significant differences in degree of starch hydrolysis across digestion time at a given number of chews (*P*<0.05).

### 3.4 SCFA production during the colonic fermentation: A comparison between 5 and 40 chews

After the INFOGEST *in vitro* digestion, the undigested rice fractions resulting from 5 and 40 chews were used as the starting materials to perform fecal batch cultures. SCFAs were then monitored during colonic fermentation for one donor over multiple days (*Fig. 4a, Fig S2,* and *Fig. S3*) and four donors (*Fig. 4b*, *4c, Fig S2,* and *Fig. S4*). Total SCFA production was significantly higher for all three rice varieties chewed 5 times compared to 40 times, regardless of whether the same donor was employed over multiple days, or four different donors were utilized during the colonic fermentations (*Fig. 4a* and *Fig. 4b*). When the number of chews was reduced from 40 times to 5 times and 4 different inocula were employed (*Fig. 4b*), the average value of total SCFAs (*Fig. 4b*) increased by 13% (basmati brown rice), 5% (pandan brown rice), and 10% (white rice).

**Fig. 4.**
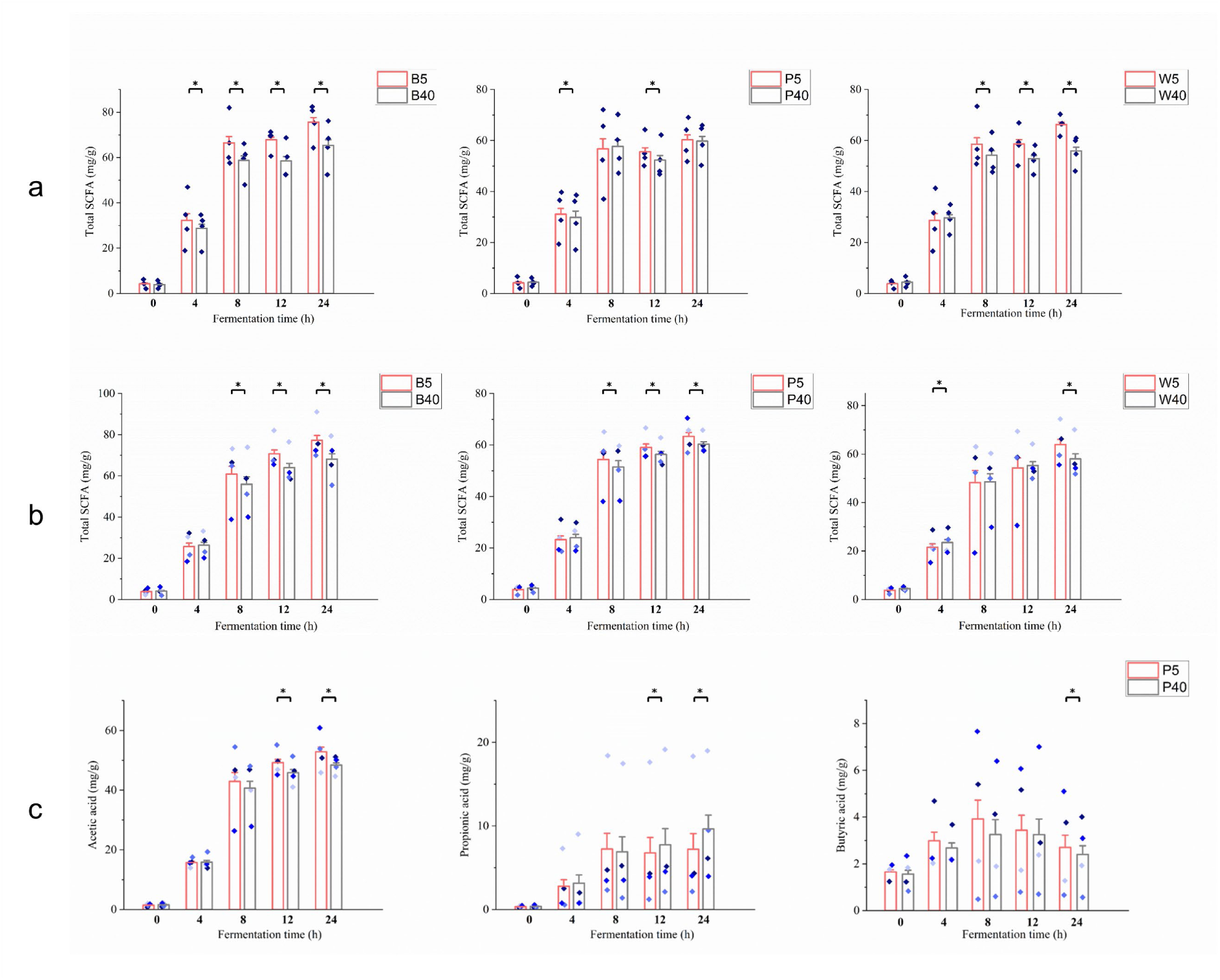
Total SCFA for one donor running the experiment in four different days (a) and four donors (n=4) (b). Acetic acid, propionic acid, and butyric acid production during *in vitro* fecal batch fermentation of pandan brown rice for four donors (n=4) (c). P5 and P40 refer to pandan brown rice samples chewed 5 and 40 times per bite. Different letters represent significant differences between 5 and 40 chews per bite for a given fermentation time (*P<*0.05).

The effect of the number of chews per bite on SCFA production was evident for acetic and propionic acids (*Fig. 4c, Fig S3, and Fig S4*). In contrast, the number of chews per bite did not impact the production of isovaleric, iso-butyric, and valeric acid (*Fig. S2, Fig. S3,* and *Fig. S4*). When microbial communities from the same donor, discovery phase, were exposed to the three undigested rice varieties that were chewed 5 and 40 times (*Fig. S3*) during the first 4 h of the colonic fermentation, there were limited differences in SCFA production across different numbers of chews per bite. After 8 h of colonic fermentation, acetic acid concentrations were significantly higher in rice varieties chewed 5 times compared to 40 times. Butyric acid showed a similar trend to acetic acid and was negatively correlated with the number of chews. However, these differences in butyric acid concentration were only significant during the first 12 hours of fermentation. The higher number of chews induced significantly more propionic acid production after 12h for all three rice varieties inoculated from the same donor on different days (*Fig S3*). When different donors were utilized in the validation phase, acetic acid and propionic acid levels, but not butyric acid production, were significantly influenced by the number of chews per bite, especially for basmati brown rice and pandan brown rice (*Fig. 4c* and *Fig. S4*). After 24 h of colonic fermentation, acetic acid concentrations were significantly higher for basmati and pandan brown rice chewed 5 times compared to 40 times. By contrast, propionic acid production was substantially lower for basmati and pandan brown chewed 5 times than 40 times.

### 3.5 Correlations between bolus properties with degree of starch hydrolysis and production of SCFA

*Fig. 5* shows a correlation heatmap reporting Spearman correlation coefficients for each comparison. The number of chews per bite was positively correlated with the number of bolus particles and total surface area (correlation coefficient: 0.488-0.878) and negatively correlated with the average size (correlation coefficient: −0.488-0.683) after the oral and gastrointestinal digestion. The correlations between the number of chews per bite with the total surface area (correlation coefficient: 0.488) and average particle size (correlation coefficient: −0.490) were weaker after *in vitro* digestion compared to the oral phase. The degree of starch hydrolysis positively correlated with the number of chews (correlation coefficient: 0.683) and was greatly influenced by the number of bolus particles and average bolus particle size. The number of bolus particles after *in vitro* digestion was negatively correlated with fiber content (correlation coefficient: −0.837), while total bolus surface area was positively correlated with fiber content (correlation coefficient: 0.717).

**Fig. 5.**
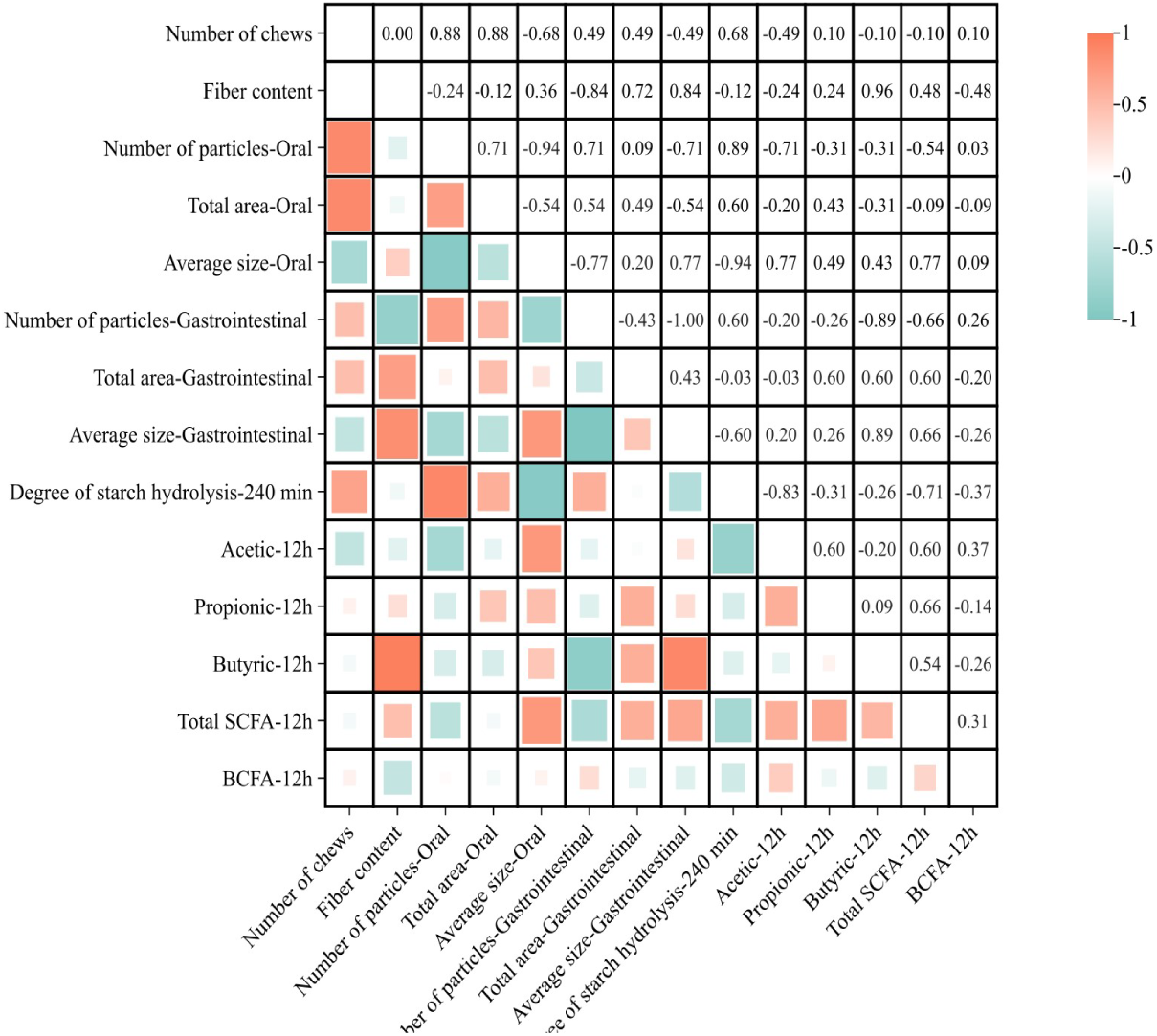
Correlations between number of chews per bite and fiber content with bolus properties after *in vivo* oral mastication and *in vitro* gastrointestinal digestion, degree of starch hydrolysis, and production of SCFAs during *in vitro* fecal batch fermentation. SCFA data used in correlation analysis was from four biological replicates (n=4).

The production of SCFA after 12 hours of fermentation was correlated with bolus particle properties, such as total surface area and average particle size. Compared with the particle properties of bolus after the oral phase, stronger correlations were found between SCFA production and particle properties after *in vitro* digestion. Acetic acid showed negative correlations with the number of chews per bite. It was highly correlated to the particle average size after oral digestion (correlation coefficient: 0.771) and the degree of starch hydrolysis after *in vitro* digestion (correlation coefficient: −0.829). Propionic acid was positively correlated with acetic acid, demonstrating a stronger correlation with total bolus surface area compared with acetic acid. Butyric acid was associated with the number of bolus particles (correlation coefficient: - 0.886) and the average size of bolus particles after gastrointestinal digestion (correlation coefficient: 0.886). Butyric acid production was strongly correlated (correlation coefficient: 0.956) with fiber content.

### 3.6 Bile salt hydrolase and microbial metabolite signatures

Bile salt hydrolase (BSH), an important enzyme involved in bile acid metabolism, was employed to demonstrate the modulation of microbial activity by bolus particle size. Higher BSH activity was observed in P5 than in P40. However, the effect was minimal in the early stages of fermentation (*Fig. 6*). After 24 hours of fermentation, higher BSH activity was found in P5 than in P40, and the differences were significant at group (*Fig. S5*) and individual levels (*Fig. 6*).

**Fig. 6.**
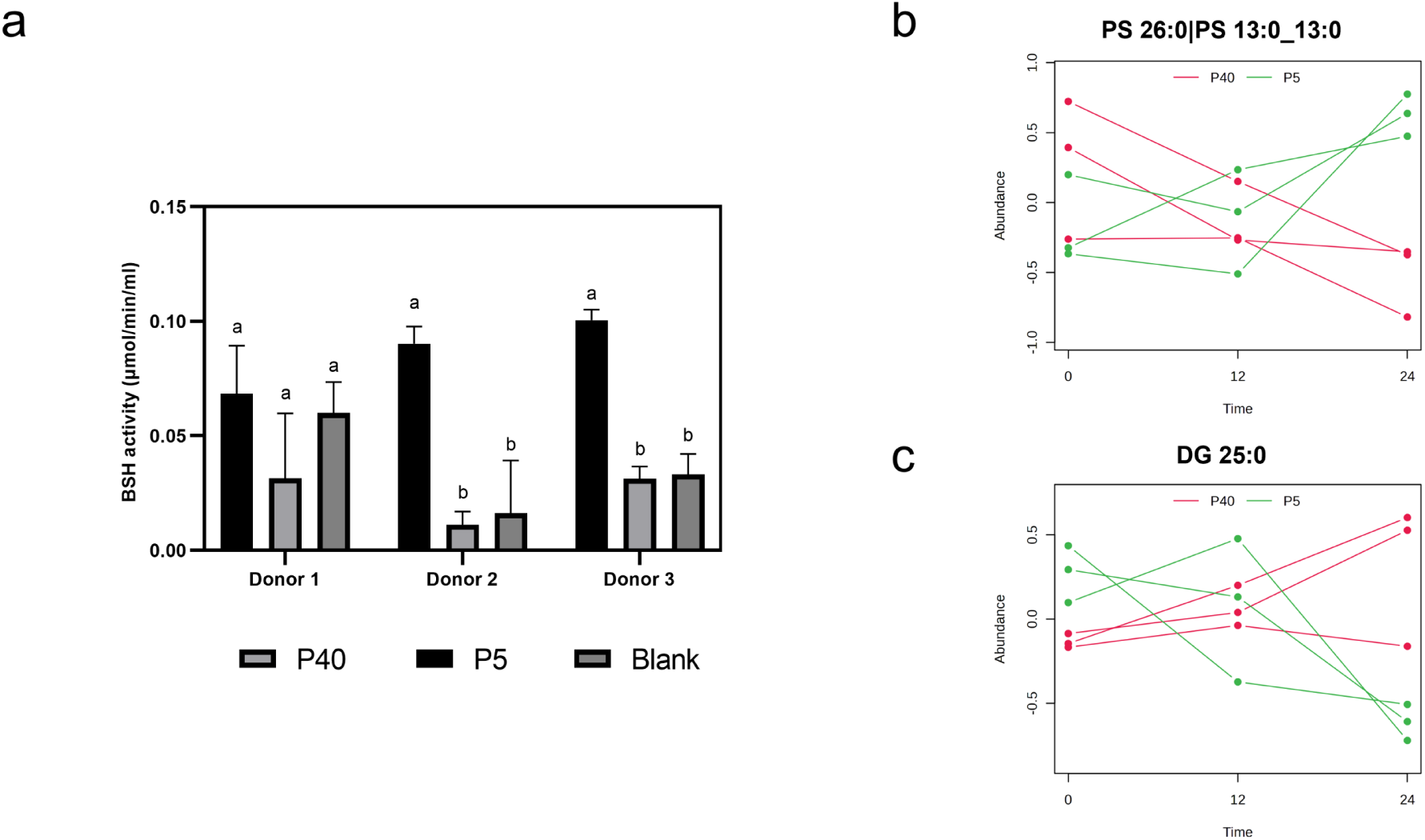
Bile salt hydrolase activity of three donors after 24 hours of *in vitro* fecal batch fermentation of pandan brown rice chewed five and forty times (a). Different letters represent significant differences between P5, P40, and fermentation blank for a specific donor (*P<*0.05). Time series + one factor and Multivariate Empirical Bayes Time-series Analysis (MEBA) analysis to explore changes in metabolites across different time points of the colonic fermentation between P5 and P40. Response of PS 26:0 (b) and DG 25:0 (c) during *in vitro* fecal batch fermentation. P5 and P40 refer to pandan brown rice samples chewed 5 and 40 times per bite.

An untargeted lipidomics approach was performed to understand further how bolus particle size modulated microbial metabolites beyond SCFA. Results of Fold Change (FC) Analysis and T-tests were combined to consider the modulation of microbiome metabolites. Metabolites with p-value < 0.05, and log_2_ FC > 2, or log_2_ FC < −2 were significantly changed by the number of chews. When the number of chews per bite increased from 5 to 40, the abundance of 103 features in the positive ionization mode and 67 in the negative mode increased. Chewing more extensively negatively affected the abundance of 27 features in positive ionization mode and 18 in negative mode after 24 hours of fermentation (*Fig. S5*). The top ten features influenced by the number of chews include sphingolipids, N-acylethanolamines, lysophosphatidylcholine, ceramides, and fatty acids (*Table S2*). Among them, two features were selected to show the effect of oral chewing behavior in modulating microbial activity and, thus, microbial metabolites throughout fermentation (*Fig. 6*). Multivariate Empirical Bayes Time-Series Analysis (MEBA) highlighted that PS 26:0 consistently increased during the fermentation of P5. At the same time, DG 25:0 decreased over time for P5, showing a significant correlation with the number of chews.

## 4. Discussion

In this study, we examined the effect of oral processing behavior on digestion and microbial metabolism by monitoring particle properties during *in vivo* mastication, *in vitro* digestion, and *in vitro* fecal batch fermentation of three rice varieties with varying fiber content. The results demonstrated that particle size is an abiotic factor influencing digestion, SCFA levels, microbial activity, and the production of other microbial lipids.

As expected, an evident influence of the number of chews per bite on bolus properties was observed in the oral phase. With increasing chewing, the cooked rice grains were disintegrated into more and smaller bolus fragments (*Fig. 1*), generating a larger total bolus surface area. This is consistent with our hypothesis and supported by previous research (Low et al., 2015; Nadia, 2022; Ranawana et al., 2010). Bolus particle size reduction in the oral phase is considered crucial for nutrient digestion (Naumova et al., 2021b), as it increases total surface area and consequently increases the accessibility of digestive enzymes (Gouseti et al., 2019). Uptake of saliva by the rice bolus increased with an increasing number of chews per bite (*Fig. S1*), which probably contributed, in addition to the oral physical breakdown, to a higher degree of starch hydrolysis, especially during the oral and gastric phase of digestion (*Fig. 3*).

*In vitro* digestion reduced bolus particle size considerably. More importantly, the impact of oral processing behavior on bolus properties persisted after the *in vitro* digestion. At the end of the *in vitro* digestion, rice chewed 40 times showed a higher number and smaller size of particles, leading to a higher total surface area than rice chewed 5 times (*Fig. 2*). Bolus particle size impacted the chemical and physical degradation of the rice particles during the digestion *in vitro* (Nadia et al., 2021b), especially during the oral and gastric phases of digestion. At the end of the intestinal digestion phase, the number of chews per bite had no significant effect on basmati rice’s degree of starch hydrolysis. In contrast, pandan and white rice showed substantial effects. Our findings suggest that the impact of oral processing behavior on starch hydrolysis decreased from the oral to the end of the intestinal digestion (*Fig. 3*). Previous research demonstrated that early glucose absorption, insulin release, and satiety responses were induced by an increased number of chews and longer oral exposure time (Goh, Chatonidi, et al., 2021), highlighting the importance of bolus properties on digestion. These differences in bolus properties after *in vitro* digestion provided a rationale to investigate further how differences in oral processing behavior and bolus properties impact gut microbiome metabolism.

During the colonic fermentation, undigested rice particles were further broken down into smaller particles (*Fig. 1* and *2*), particularly for rice chewed 40 times. To prove that particle size is a key abiotic factor regulating the gut microbiome, two *in vitro* fecal batch fermentation studies were performed: one donor repeated multiple times (n=4) for the discovery phase and different donors (n=4) for the validation. In the discovery phase, fewer chews per bite (5 times) promoted the production of acetic acid and butyric acid but decreased propionic acid production compared to 40 chews per bite. This indicates that chewing, thus bolus particles, might chronically impact the production of SCFAs. Secondly, more biological replicates were included to investigate inter-individual variability *in vitro*. Acetic acid concentration was higher after 5 chews per bite, and propionic acid was boosted with 40 chews per bite. When examining specific SCFAs, a larger surface area was positively correlated with propionic acid and negatively correlated with acetic acid. Interestingly, acetic acid and propionic were positively correlated (*Fig. 5*), despite their opposite correlation with surface area. Macfarlane and co-authors found that adherent species are considerably less efficient than nonadherent bacteria during the fermentation of starch and produce less acetic acid and butyric acid (Macfarlane & Macfarlane, 2006). This partly explains our results and indicates that a higher surface area facilitates more production of propionic acid through the 1,2- propanediol pathway rather than the succinate and acrylate pathways (Oliphant & Allen-Vercoe, 2019). On the other hand, butyric acid production was positively correlated with the particle size after digestion. This is consistent with a previous study (Tuncil et al., 2018). We speculated that the substrate and the ability of microorganisms to adhere to food particles (Ze et al., 2013) could rule out this symbiotic relationship, where primary degraders attach to particles and select microorganisms that favor the release of specific microbial metabolites (Wang et al., 2019). Ultimately, when we tested our second hypothesis, we found that the undigested rice derived from 5 chews, thus larger particles and smaller surface area compared to 40 chews, boosted the total SCFA production (*Fig. 4*). This was inconsistent with our hypothesis but is aligned with a recent *in vivo* study (Letourneau et al., 2024). This research study found low levels of SCFAs in small fecal particle sizes and a correlation between small fecal particle sizes and long transit times, suggesting that SCFAs may have been absorbed by enterocytes. However, in our static *in vitro* system that does not currently incorporate a cell compartment regulating absorption, we also noticed that smaller particle sizes were associated with lower levels of SCFAs.

Apart from particle size and surface area, starch hydrolysis caused by chewing might also contribute to SCFA production, especially acetic acid, which was strongly correlated with the degree of starch hydrolysis (*Fig. 5*). Rice is a food rich in starch, which is composed of two types of polysaccharides, amylose and amylopectin (Martens et al., 2018). Several research works have reported that compared with amylose, amylopectin is easier to hydrolyze during gastrointestinal digestion and to be used by gut microbiota (Pan et al., 2023; Z. Zhou et al., 2013). In our study, extensive chewing resulted in a higher degree of starch hydrolysis, which may reduce amylopectin levels and increase the availability of amylose, which hydrolyzes more slowly. This may explain why acetic acid, an early intermediate metabolite, was produced at higher concentrations when there was a lower degree of starch hydrolysis on the sample with limited chewing. However, it is essential to point out that the impact of bolus particle size on digestion and microbial fermentation might partly depend on rice variety. In our study, white rice exhibited more starch hydrolysis, a greater number of particles, and smaller particle sizes after *in vivo* chewing and *in vitro* digestion compared to pandan and basmati rice, irrespective of the number of chews. We suspect that the higher glycaemic response to white rice may be in part attributed to its bolus properties and the associated increased starch hydrolysis due to lower fiber content, greater saliva uptake, and smaller bolus particles (Liu et al., 2021).

Our results indicate that the gut microbiome responds differently to particle sizes rather than to different substrate compositions. The correlation analysis showed that the type of rice only influenced butyric acid production. The literature lacks consensus, linking outcomes with particle size (Thakkar et al., 2020) and substrates (Yao et al., 2023). However, our results aligned with a recent study that embraced the complexity of the entire gastrointestinal tract and showed that intensive chewing of rice reduced total SCFA production and increased propionic acid concentration *in vitro* (Kim et al., 2022). This study showed that SCFA production was significantly attributed to particle size and surface area differences. This led us to hypothesize that particle size and surface area may influence other microbial metabolites through specific enzymatic activities (Mohammed & Guda, 2015). In the field of soil microbiome, it has been reported that microbial enzymatic activity, such as phosphatase and peroxidase, differed across particles of different sizes (Lagomarsino et al., 2009; Zhang et al., 2015). Thus, we measured BSH activity to demonstrate that particle size may have influenced microbial enzymatic activity. Our results highlighted that increased chewing per bite led to lower BSH enzyme activity. This enzyme deconjugates primary bile acids, releasing taurine and glycine. Thus, it is tempting to speculate that a higher BSH activity observed in less chewed samples could protect commensal bacteria from bile acid toxicity, contribute to bacterial survival and colonization, and confer a nutritional advantage since released amino acids might be used as nitrogen and energy sources. Overall, this indicates that particle size altered the BSH levels to adapt to different particle sizes and areas, possibly resulting in changes in the metabolite pool.

To confirm this, we carried out an untargeted lipidomics approach. We observed that the employed substrates containing different particle sizes significantly affected the abundance of more than 150 tentatively identified features (*Fig. S5*), indicating that particle size influenced the signature of microbial metabolites. For example, phosphatidylserine (PS) and diacylglycerol (DG) are metabolic intermediates during bacterial membrane formation (*Fig. 6*). The differences observed in our study may suggest that particle size promoted the activity of specific microbial communities (Ryan et al., 2023). *Table S2* summarizes the top 10 metabolites, showing the impact of food particles on microbial activity. Among the 10 top metabolites, sphingolipids (SLs) were the most abundant, and they could be synthesized by *Bacteroidetes* and certain *α-Proteobacteria* species by the enzyme serine palmitoyltransferase (Brown et al., 2019). The differences in SLs and other lipids, such as fatty acids, further support the idea that the gut microbiome adapts its microbial activity to particle size, resulting in different metabolite signatures.

This study provides valuable insights into how bolus particle size affects digestion and the function of the gut microbiome. However, there are some limitations to consider. First, the results of the present study were from *in vitro* systems with limited complexity compared with the human body. Gastric emptying, dynamic digestion, and nutrient absorption were not considered during the *in vitro* digestion (Mackie et al., 2020; Woolnough et al., 2008). Metabolites accumulate during *in vitro* fermentation and may affect microbial activity (den Besten et al., 2013). The gut microbiome and metabolite responses to oral chewing behavior observed in rice may differ from other foods. Further *in vitro* and *in vivo* studies are necessary before conclusive evidence can be obtained regarding the impact of oral chewing behavior on the gut microbiome. Despite these limitations, our findings build on a growing body of research demonstrating bolus particle size’s influence on digestive kinetics, metabolism, and microbiome function, which are not exclusively driven by dietary patterns but also by the degree of physical and chemical degradation after gastrointestinal digestion. Further research is needed to understand whether oral processing behavior chronically impacts digestion and the functionality of the gut microbiome, modulating enzymatic activities, such as BSH, and the pattern of microbial metabolites like SCFAs that might affect the intestines and peripheric organs.

## 5. Conclusions

The effect of oral processing behavior on the bolus properties of three rice varieties persisted from *in vivo* mastication to *in vitro* digestion and colonic fermentation. Small bolus particle sizes and large total bolus surface area induced by more chewing increased the degree of starch hydrolysis during gastrointestinal digestion and, in turn, influenced microbiome enzymatic activity and metabolite production by the gut microbiota.

Our study highlighted that gut microbiome adjusted their microbial activity, such as BSH activity and SCFA production, to different bolus particle sizes. *In vivo*, these metabolite differences may further influence human metabolism in many ways. Our results suggest that consuming rice with fewer chews may provide more energy for the human gut microbiota, as we observed increased production of total SCFAs with less chewed rice. Increased chews per bite might contribute to earlier satiation by, amongst other factors, higher propionic acid production. Chewing behavior, thus bolus particles, regulates other metabolites beyond SCFA that may exert local effects on intestinal epithelial cells.

We conclude that (a) differences in rice bolus properties caused by differences in chewing behavior are maintained throughout *in vitro* gastrointestinal digestion and colonic fermentation and (b) chewing behavior and bolus properties of rice influence carbohydrate digestion and microbiome functionality during the colonic fermentation that might regulate human metabolism. Ultimately, future research should validate these findings using other foods and explore whether the gut microbiome composition and its functionality are modulated by oral processing behavior in dedicated dietary interventions. As a key abiotic factor, food particles can provide innovative strategies for effectively treating digestive disorders linked to the gut microbiota by directly altering the function of the gut microbiome.

## CRediT authorship contribution statement

**Zhen Liu:** Investigation, Methodology, Data curation, Formal analysis, Writing – original draft. **Ciarán Forde:** Conceptualization, Methodology, Writing – review & editing. **Markus Stieger:** Conceptualization, Methodology, Writing – review & editing. **Josep Rubert:** Conceptualization, Methodology, Writing – original draft, Writing – review & editing.

## Acknowledgments

Zhen Liu received a PhD scholarship from the China Scholarship Council (CSC No. 202206850004). Core funding from the Food Quality and Design and Sensory Science and Eating Behaviour groups and the Dutch Top-Consortium for Knowledge and Innovation Agri & Food (TKI-Agri-food) Project Restructure (TKI 22·150) supported this research throughout different stages.

## Data availability

The datasets generated during the current study are deposited in the Zenodo repository under the DOI https://doi.org/10.5281/zenodo.14849523

