## Supplemental figures and tables for "Chewing behavior and bolus particle size of rice shape gut microbiota functionality and microbial metabolite signatures"

**Supplementary data**

List of Supplementary Tables:

**Table. S1** Composition (water, carbohydrate, fiber, and protein content) of three kinds of rice.

**Table. S2** Top 10 features regulated by the number of chews in colonic fermentation. The features were selected according to Fold Change (FC) Analysis and T-tests. The feature ID is defined as "retention time (min)" / "m/z". It includes the name of the compound tentatively identified, the molecular formula, the INCHIKEY, and ontology.

List of Supplementary Figures:

**Fig. S1** Saliva uptake (%) of three rice varieties after different numbers of chews. Different capital letters denote significant differences in the rice variety's number of chews per bite. Different lowercase letters indicate significant differences across rice varieties within the number of chews per bite ($P<0.05$). Error bars represent standard deviation.

**Fig. S2** BCFA of one donor over multiple days (a) and multiple donors (b). B5 and B40 refer to basmati brown rice samples chewed 5 and 40 times. P5 and P40 refer to pandan brown rice samples chewed 5 and 40 times. W5 and W40 refer to white rice samples chewed 5 and 40 times. Dots with the same colors show the values of one donor.

**Fig. S3** Acetic acid, propionic acid, and butyric acid production during *in vitro* fecal batch fermentation of basmati brown rice (a), pandan brown rice (b), and white rice (c) of one donor over multiple days. Isovaleric, isobutyric acid, and valeric acid production during *in vitro* fecal batch fermentation of basmati brown rice (d), pandan brown rice (e), and white rice (f) of one donor in multiple days. B5 and B40 refer to basmati brown rice samples chewed 5 and 40 times. P5 and P40 refer to pandan brown rice samples chewed 5 and 40 times. W5 and W40 refer to white rice samples chewed 5 and 40 times. Dots with the same colors show the values of one donor.

**Fig. S4** Acetic acid, propionic acid, and butyric acid production during *in vitro* fecal batch fermentation of basmati brown rice (a), and white rice (b) for four donors. Iso valeric, iso butyric acid, and valeric acid production during *in vitro* fecal batch fermentation of basmati brown rice (c), and white rice (d) of four donors. B5 and B40 refer to basmati brown rice samples chewed 5 and 40 times. W5 and W40 refer to white rice samples chewed 5 and 40 times. Dots with the same colors show the values of one donor.

**Fig. S5** Bile salt hydrolase activity after 24h of fermentation of pandan brown rice (a). Different letters represent significant differences between P5, P40, and fermentation blank at a given fermentation time ($P$<0.05). Principal Component Analysis (PCA) plots based on the levels of microbial metabolites under ESI positive (b) and ESI negative (c) after 24 hours of *in vitro* fermentation. Volcano plots for enriched metabolites in P40 or P5 samples after 24 hours of *in vitro* fecal batch fermentation under ESI positive mode (b) and ESI negative mode (c). The x-axis represents $\log_2$ fold change (FC), and the y-axis represents $-\log_{10}$ p-value. Metabolites with significant changes (p-value < 0.05, log2 fold change > 2) were considered upregulated in P40 compared with P5. Metabolites with significant changes (p-value < 0.05, log2 fold change < -2) were considered downregulated in P40 compared with P5. The top 5 features regulated by the number of chews in colonic fermentation were highlighted in the graph, which was selected according to FC and p-value. P5 and P40 refer to pandan brown rice samples chewed 5 and 40 times per bite.

| | Water (g/100g ) | Carbohydrate (g/100 g) | Fiber (g/100 g) | Protein (g/100 g) |
|---|---|---|---|---|
| Basmati brown rice | 13.50 | 69.00 | 6.10 | 8.40 |
| Pandan brown rice | 12.50 | 66.00 | 8.50 | 9.50 |
| White rice | 12.50 | 78.00 | 1.50 | 7.00 |

Table. S1

| Feature ID (RT / m/z) | Compound name | Group marker | Molecular formula | InChIKey | Polarity | Ontology |
|---|---|---|---|---|---|---|
| 4.78/566.3482 | SL 31:6;O2\|SL 13:1;O/18:5;O | P40 | C31H51NO6S | CDNSESAZPBRHIV-UHFFFAOYSA-N | ESI (+) | SL |
| 9.01/430.3771 | NAE 26:5 | P40 | C28H47NO2 | SKLYJTADGQXJFC-UHFFFAOYSA-N | ESI (+) | NAE |
| 7.93/637.5232 | SL 34:0;O2\|SL 12:0;O/22:0;O | P40 | C34H69NO6S | XTDFIBNYURUOID-UHFFFAOYSA-N | ESI (+) | SL |
| 5.26/581.3528 | SL 31:7;O2\|SL 13:2;O/18:5;O | P5 | C31H49NO6S | SIXSMEBMZCATFH-UHFFFAOYSA-N | ESI (+) | SL |
| 3.29/562.4397 | Cer 34:5;O4\|Cer 12:1;O3/22:4(2OH) | P5 | C34H59NO5 | BTYFLZADABMLMD-UHFFFAOYSA-N | ESI (+) | Cer |
| 0.78/600.3849 | Cer 34:8;O4\|Cer 12:2;O3/22:6(2OH) | P40 | C34H53NO5 | BOKCMWPMOUIEQW-UHFFFAOYSA-N | ESI (-) | Cer |
| 4.36/680.4379 | DMPE 30:5\|DMPE 8:0_22:5 | P40 | C37H64NO8P | SNMQXHILMFBXMI-UHFFFAOYSA-N | ESI (-) | DMPE |
| 2.51/731.374 | SMGDG 27:4\|SMGDG 8:0_19:4 | P40 | C36H60O13S | XCHGRSOIXPBXMI-UHFFFAOYSA-N | ESI (-) | SMGDG |
| 7.07/291.1998 | FA 18:4;O | P5 | C18H28O3 | JZPSRJYCHQSSRG-UHFFFAOYSA-N | ESI (-) | FA |
| 3.87/456.2331 | LPC 10:0 | P5 | C18H38NO7P | SECPDKKEUKDCPG-UHFFFAOYSA-N | ESI (-) | LPC |

Table. S2

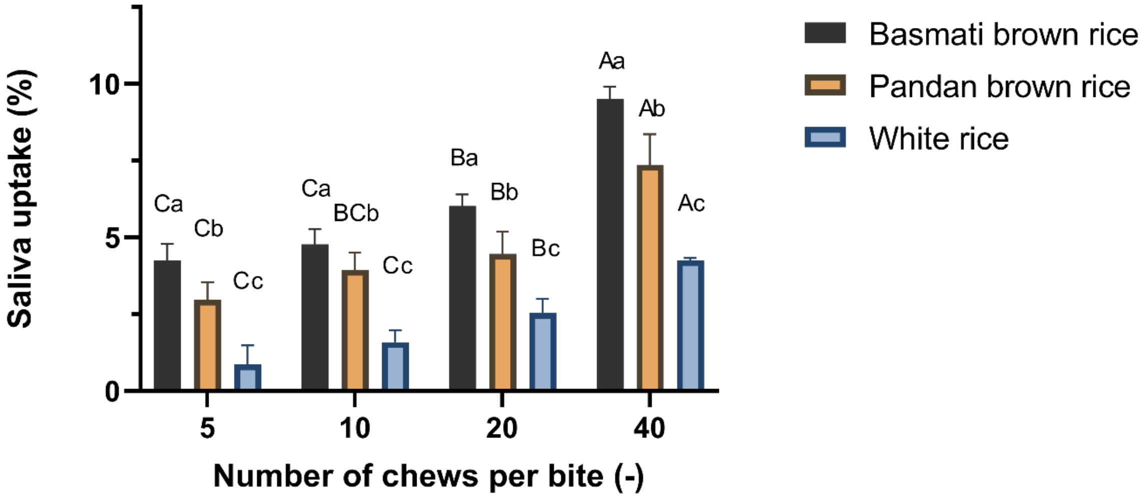

Saliva uptake (%)
0
5
10
Number of chews per bite (-)
5
10
20
40
Ca
Cb
Cc
Ca
BCb
Cc
Ba
Bb
Bc
Aa
Ab
Ac
Basmati brown rice
Pandan brown rice
White rice


Fig. S1

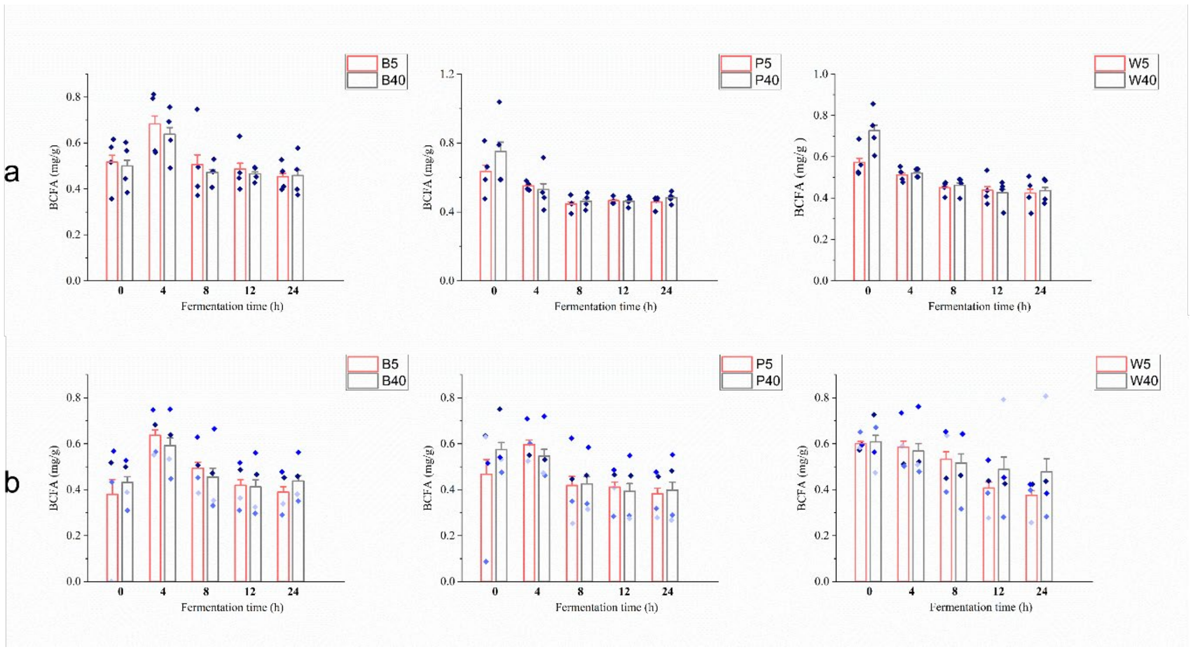


Fig. S2

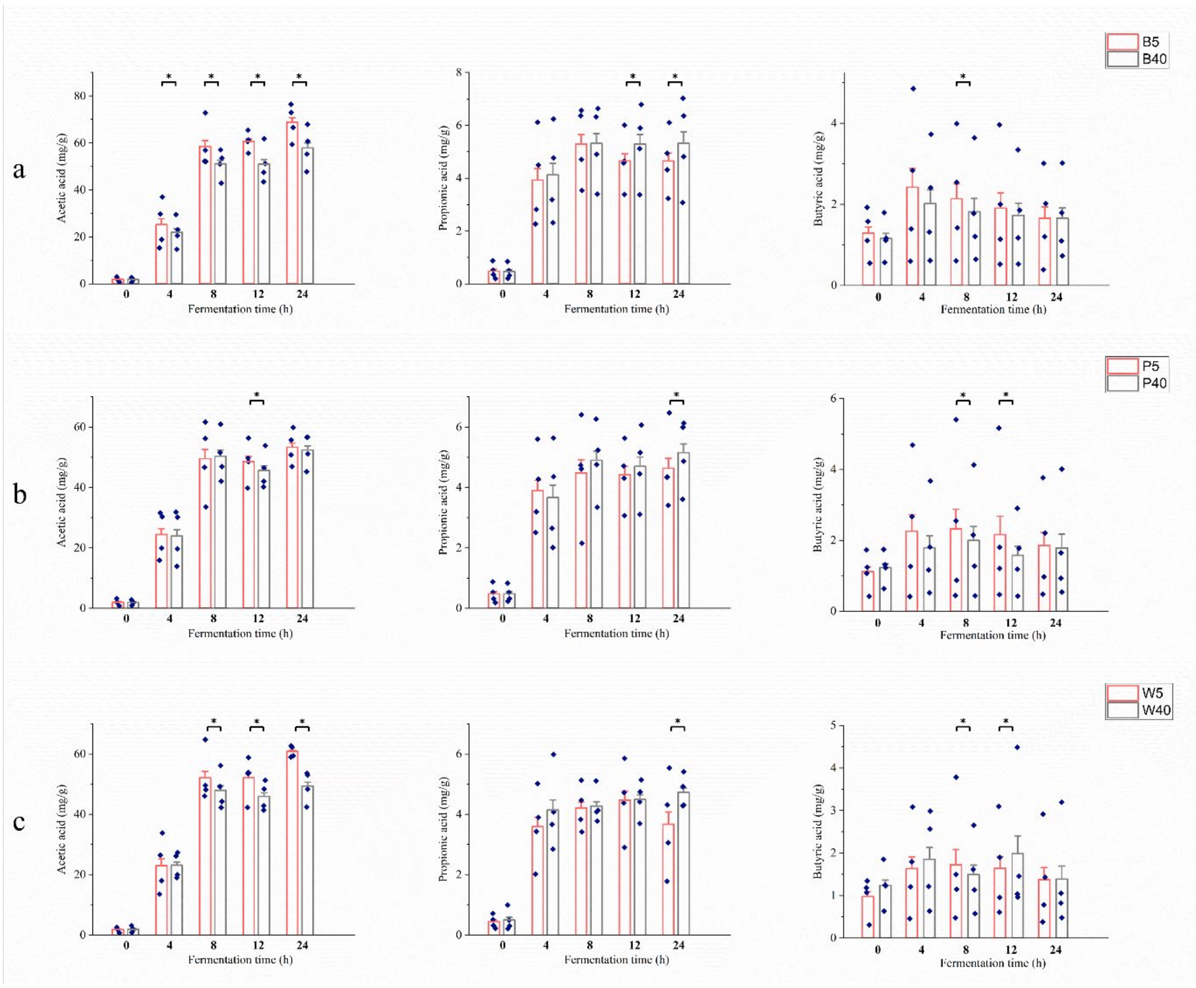
a
b
c
B5
B40
P5
P40
W5
W40
Acetic acid (mg/g)
Propionic acid (mg/g)
Butyric acid (mg/g)
Fermentation time (h)

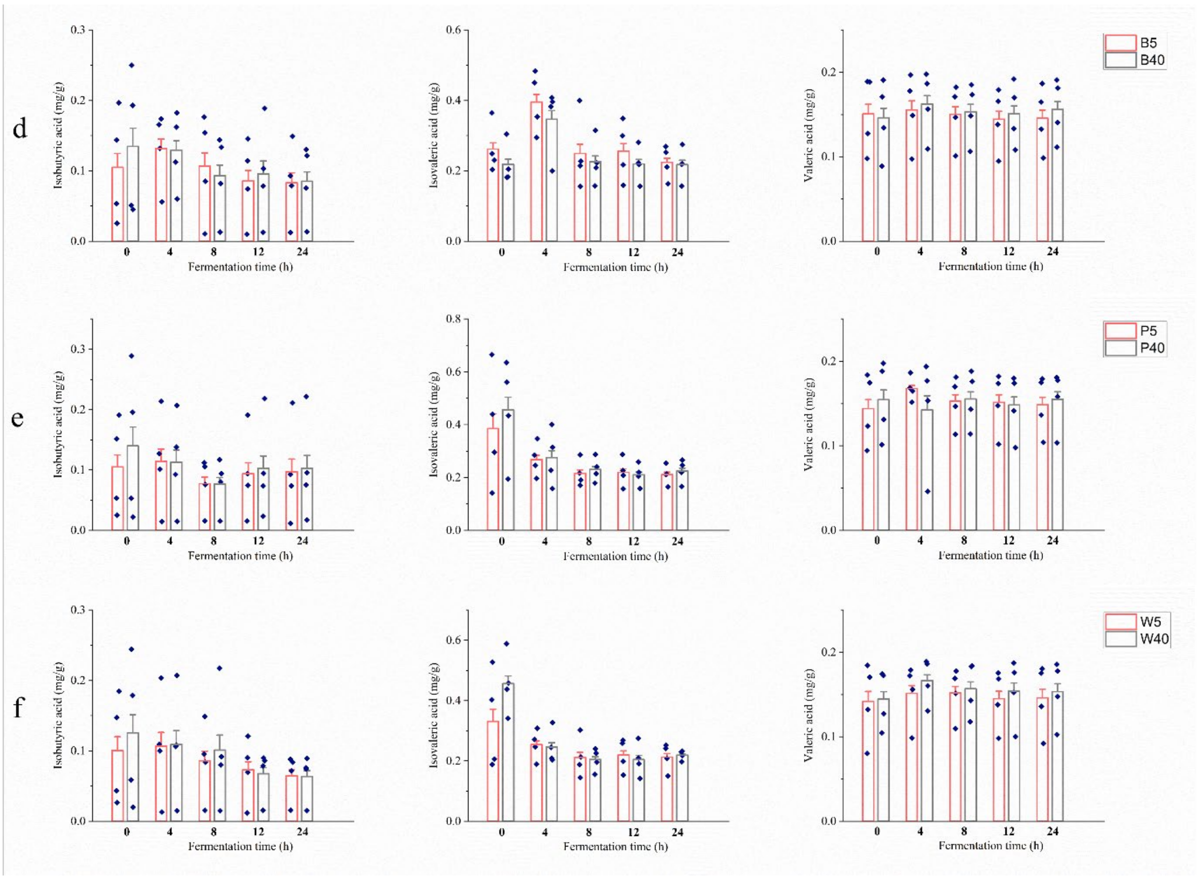
d
e
f
B5
B40
P5
P40
W5
W40
Isobutyric acid (mg/g)
Isovaleric acid (mg/g)
Valeric acid (mg/g)
Fermentation time (h)

Fig. S3

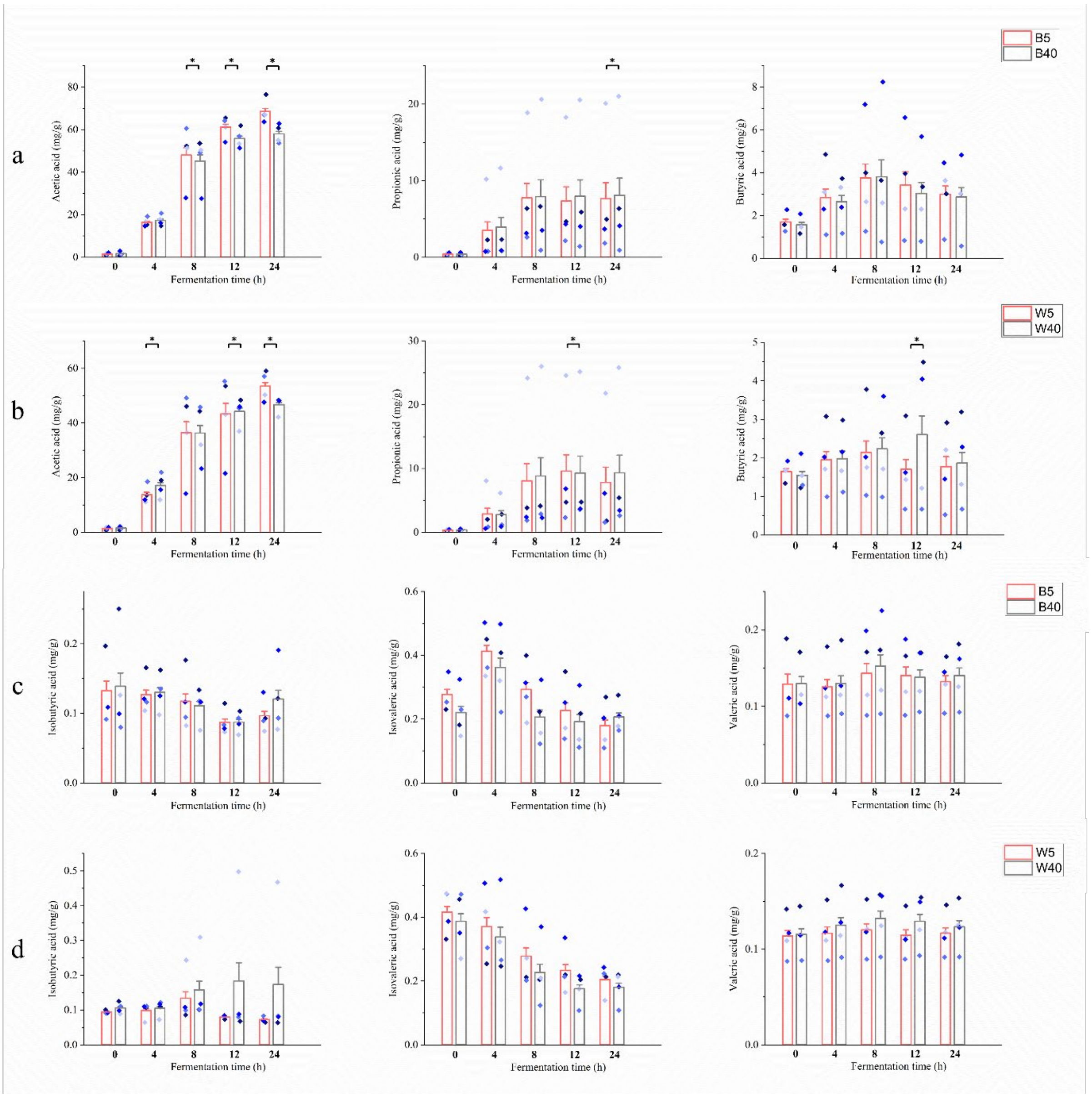
B5
B40
W5
W40
a
b
c
d
Acetic acid (mg/g)
Propionic acid (mg/g)
Butyric acid (mg/g)
Isobutyric acid (mg/g)
Isovaleric acid (mg/g)
Valeric acid (mg/g)
Fermentation time (h)

Fig. S4

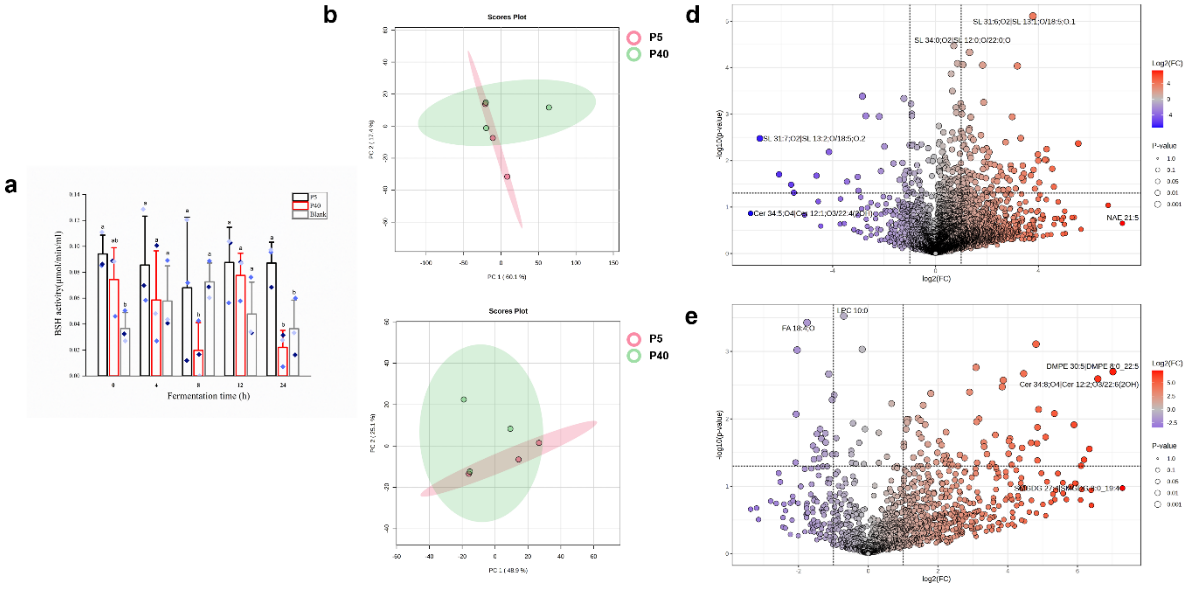
a
b
d
e
Scores Plot
Scores Plot
P5
P40
P5
P40
BSH activity(μmol/min/ml)
Fermentation time (h)
P5
P40
Blank
log2(FC)
-log10(p-value)
Log2(FC)
P-value
log2(FC)
-log10(p-value)
Log2(FC)
P-value

Figure. S5
